# Pediatric Hodgkin disease in Brazil: a disease with a highly heterogeneous treatment

**DOI:** 10.1101/628735

**Authors:** Fernanda Lagares Xavier Peres, Antônia Pardo Chagas, Natália Dassi, Thais G. Almeida, Ângela Rech Cagol, Cecília Fernandes Lorea, Pablo Santiago, Laura Garcia de Borba, Liane Esteves Daudt, Mariana Bohns Michalowski

**Affiliations:** Medical Student, Universidade Federal de Ciências da Saúde de Porto Alegre (UFCSPA); Medical Student, Universidade Federal do Rio Grande do Sul (UFRGS); MD, PhD, Universidade de Caxias do Sul (UCS); MD, Universidade Federal de Pelotas (UFPel); MD, Hospital São Vicente de Paula (HSVP); MD, Hospital da Criança Santo Antônio (HCSA); MD, PHD, Hospital de Clínicas de Porto Alegre (HCPA), Universidade Federal do Rio Grande do Sul (UFRGS)

**Author notes:** **Corresponding Author:** Profa. Mariana Bohns Michalowski, Serviço de Oncologia Pediátrica, Hospital de Clínicas de Porto Alegre, Rua Ramiro Barcelos, 2350, CEP 90035-903, Porto Alegre – RS.

## Abstract

Standard of care and protocols for the treatment of pediatric cancer lead to a clear improvement in survival rates and quality of life. Little is known about how these treatments are implemented in Brazil. Our study aimed to evaluate children treated for Hodgkin disease (HD) in south Brazil between 2002 and 2013 through the analysis of medical records in 6 different centers.

**Results:** Fifty-nine children and adolescents were included. The median age was 12 years (range 3-18 years). Male:female ratio was 1.95:1. Localized disease (stage I/II) was observed in 30 patients (50.8%) while the remaining 29 (49.2%) had advanced disease (Stage III/IV).

The chemotherapeutic treatment schema was different among services and comprised three different based protocols. ABVD schema was the most frequently used (52 children (88.1%)). The number of cycles was highly variable (4-16 cycles) even at the same clinical stage and with similar clinical response.

**Conclusion:** These data highlight the importance of turning the “best practice policies” readily available to all pediatric oncologists. Local protocols allow integrative studies among centers that would certainly maintain or improve cure rates, reduce long-term toxicity and evaluate specific biological characteristics of these diseases in our population. For these reasons, we reinforce the idea that standardization of treatment in pediatric oncology is a child health priority and also a viable low-cost strategy to improve care in middle-income countries such as Brazil.

## INTRODUCTION

Many times, improvement in healthcare focuses in the use of new drugs. In fact, in many situations this is the most reliable and important measure available. Recently, in pediatric oncology there was an increase in the use of immunotherapy as a therapeutic choice in different diseases. Although they are very effective, their prices are frequently high which makes them a significant spent in middle income countries that have many other essentials health issues to be addressed. To rationalize the use of these new agents and indicate them correctly, one must be sure that an effective first-line treatment is being used. It is also important to notice that through the years the Brazilian population benefits of a unified health system (SUS) that allows access to most treatment. In this reality, children with cancer can have access to Pediatric Oncology Centers through a local network. Unfortunately, data about treatment, evolution and long-term effects are not routinely recorded.

In this context, we looked for one important marker of quality of care in pediatric oncology assistance: the use of protocols or, in their absence, clinical practice guidelines. Although there is no national guideline, Hodgkin disease (HD) is a neoplasia that has a standard of care with chemotherapeutic agents available in Brazil and good international results. In our study, we were able to conclude that there was a high inter-institutional heterogeneity of treatment but also there was not a standard treatment inside each institution.

## Methods

This is a descriptive retrospective study based on medical records and the following data were analyzed: age at diagnosis, sex, stage, presence of Bulky disease at diagnosis, therapeutic regimen, first treatment response, relapse, high dose chemotherapy regimen and radiotherapy. 59 children and adolescents registered in six different pediatric oncology centers in South Brazil between 2002 and 2013, with clinical and histopathological diagnosis of Hodgkin disease (HD) were included in this study. The study followed institutional ethical guidelines.

Patients were staged according Ann Arbor staging criteria and the presence of B symptoms (fever greater than 38°C for at least 3 consecutive days, night sweats and unexplained weight loss greater than 10% of body weight within the last 6 months). The procedures used for staging were clinical history, physical exam, CT scans (neck, chest and abdomen), gallium scintigraphy and bone marrow biopsy.

The schema of chemotherapy and radiotherapy received was registered, as well as response and status.

## Results

Fifty-nine children and adolescents registered in six different pediatric oncology centers in South Brazil with diagnosis of HD were included. The median age was 12 years (range 3-18 years). Male:female ratio was 1.95:1. Localized disease (stage I/II) was observed in 30 patients (50.8%) while the remaining 29 (49.2%) had advanced disease (Stage III/IV).

The chemotherapeutic treatment schema was different among services and comprised three different based protocols: ABVD (doxorubicin, bleomycin, vinblastine and daunorubicin) in 52 children (88.1%), BEACOPP (bleomycin, etoposide, doxorubicin, cyclophosphamide, vincristine, procarbazine and prednisone) in 5 (8.5%) and Euronet (vincristine, adriamycin, etoposide and prednisone) in 2 (3.4%). The number of cycles was highly variable. Children treated with ABVD received from 4 up to 16 cycles of treatment even at the same clinical stage and with similar clinical evolution (Table 1).

**Tabela I.** Presentation, treatment and evolution of patients

| Clinical Presentation |  |  |  |  |  | First-line treatment |  |  | Evolution |  |
| --- | --- | --- | --- | --- | --- | --- | --- | --- | --- | --- |
| Ann Harbor | N | B symptoms (n) | Median age (years) | Sex | Bulky disease (n) | Chemotherapy x cycles (n) | Radiotherapy (n) | Response (n) | Relapse(n) | Last information(n) |
| 1 | 8 | 0 | 10,5 | Male 6<br>Female 2 | Nenhum | ABVD x 4 (5)<br>ABVD x 6 (2)<br>Euronet (1) | 3 | CR = 8 | 1 | RC = 8 |
| 2 | 22 | 11 | 12,5 | Male 15<br>Female 7 | 8 | ABVD x 4 (6)<br>ABVD x 6 (10)<br>ABVD x 8 (3)<br>R-ABVD x 4 (1)<br>BEACOPP (1) | 13 | CR = 16<br>PR = 3<br>PD = 1<br>NI= 2 | 7 | CR = 19<br>On treatment= 3 |
| 3 | 22 | 11 | 13 | Male 13<br>Female 9 | 8 | ABVD x 3 (1)<br>ABVD x 4 (6)<br>ABVD x 6 (3)<br>ABVD x 8 (8)<br>ABVD x 16 (1)<br>BEACOPP (2)<br>Não informado (1) | 16 | CR = 14<br>PR = 4<br>PD = 2<br>NI= 2 | 4 | CR = 18<br>On treatment= 2<br>Death = 2 |
| 4 | 7 | 5 | 13 | Male 5<br>Female 2 | 2 | ABVD x 6 (3)<br>ABVD x 7 (1)<br>ABVD x 8 (2)<br>Euronet (1) | 4 | CR = 5<br>PR = 1<br>SD = 1 | 2 | CR = 6<br>Death = 1 |

NI: Not informed; CR: complete response; PR: partial response; PD: progressive disease; SD: stable disease.

Radiation therapy was used in 34 patients (58.6%). As described in Table 1, this indication did not follow stage or remission criteria. In this cohort three (5%) patients died, one of treatment related toxicity and 2 of disease progression.

## DISCUSSION

HL is a paradigm in pediatric oncology since the goal of treatment is not only to heal but also to guarantee quality of life and social insertion for these children. Through our study, we were able to conclude that treatment in our reality was highly heterogeneous among institutions but also that there was not a standard of care even inside each institution ^1^. Our study represents a small sample of children with a specific disease (HD) from only one state in Brazil and did not evaluate the use of clinical and radiological criteria of stage and response. We can assume that discrepancies are even greater among different regions of our country with lower socio-cultural status or less access to health system and that they are probably not restricted to HD.

It is widely described that patients that are enrolled and followed up in a protocol of treatment have better prognosis and probably less adverse effects than those who are not ^2^. We also know that networking is fundamental to facilitate access to exams and research, but also to allow a better comprehension of the local reality ^3^. Although we have shown in our study that survival rate in our population is similar to those described in international studies, we can infer from these data that some children received an excessive treatment leading to a possible long-term toxicity.

These data highlight the importance of turning the “best practice policies” readily available to all pediatric oncologists. Local protocols allow integrative studies among centers that would certainly maintain or improve cure rates, reduce long-term toxicity and evaluate specific biological characteristics of neoplasias in our population. For these reasons, we reinforce the idea that standardization of treatment in pediatric oncology is a child health priority and a viable low-cost strategy to improve care in middle-income countries such as Brazil ^4,5^.

